# Contentopic mapping in ventral and dorsal association cortex: the topographical organization of manipulable object information

**DOI:** 10.1101/2023.11.29.568856

**Authors:** J. Almeida, S. Kristensen, Z. Tal, A. Fracasso

## Abstract

Understanding how object information is neurally organized is fundamental to unravel object recognition^1–4^. The best-known neural organizational principle of information is topographical mapping of specific dimensions. Such maps have been shown mainly for sensorimotor information within sensorimotor cortices (e.g., retinotopy)^5–9^. Thus, here we ask whether there are topographic maps – by analogy, contentopic maps – for mid-level object-related dimensions. We used functional magnetic resonance imaging and population receptive field analysis^7^ to measure tuning of neural populations to selected manipulable object-related dimensions. Here we show maps in dorsal and ventral occipital cortex that code for the score of each object on each target dimension in a linear progression following a particular direction along the cortical surface. Maps for each dimension are distinct, and consistent across individuals. Thus, object information is coded in multiple topographical maps – i.e., contentopic maps. These contentopic maps refer to intermediate level visual and visuomotor representations, and are potentially computed from the grouping of lower-level information through non-linear transformation following gestalt principles^4,10^. This shows that topography is a widespread and non-incidental strategy for the organization of information in the brain that leads to greatly reduced connectivity-related metabolic costs and fast and efficient readouts of information for stimuli discrimination^11,12^.

---

A significant part of the life of an animal is spent on recognizing and identifying what is present in the environment in order to make informed decisions on essential issues such as foraging, mating, fighting, or fleeing. Object recognition became even more central for humans because our environment evolved beyond the natural world, and a myriad of human-made objects invaded our daily routines. A starting point for understanding how we recognize objects is focusing on how object-specific information is represented and organized in the brain^1–4^.

The best-known organizational principles of information in the brain are those present at primary sensorimotor cortices. For instance, visual cortex is organized by cortical columns that locally and continuously code for orientation of the visual stimulus^5^. Moreover, individual cortical columns refer to particular locations in the visual field, with proximity in neural space mirroring proximity in the visual field. This organization musters a continuous and topographical map of the visual field across the different cortical columns that gradually follows retinal/visual field location of the stimulus (i.e., a retinotopic map^6,7^, for similar topographical maps in other sensorimotor cortices see^8,9^).

Importantly, extant data on mid- and high-level object processing may point to a similar pattern of organization of object information in the brain. Work on non-human primates seems suggestive of a columnar organization within association cortex that is responsive to mid-level object-specific features such as complex shapes or combination of shapes, or object textures^13^. Moreover, adjacent columns code for ever so slightly different mid-level features^13^, suggesting similar topographical properties to those present in the columns within early visual cortex. Interestingly, recent work on object-driven human neural responses within association cortex showed that these regions code for object-related dimensions^1,14–21^. Examples of the dimensions tested include, among others, real object size^15^, shape curvature and spikiness^17,18,20^, or visual field preferences and eye fixation patterns on the processing of different object domains^16,19,21^. These attempts have revealed very large organizational mosaics and extremely coarse topographies, whereby different regions dedicated to different dimensions or categories have been proposed (e.g., regions that code for large objects that are physically separated from regions that code for small objects^15^). It is not clear whether this level of explanation and degree of topography is sufficient to understand the organization of object knowledge in the brain, or whether we need to inspect finer-grained organizational levels and look for true topographical maps – i.e., look for finer-grain continuous mapping of cortical preferences within the cortical surface as a function of the (continuous) levels of an object-related dimension.

One aspect that may have deterred finding such topographic maps relates to the nature and precise content of object-related dimensions, which are considerably more elusive than those of the sensorimotor dimensions that rule the organization of sensorimotor cortices. The complexity of the world dictates that contrary to tone frequency of a sound, or the eccentricity of a visually presented stimulus in the visual field, an individual object-related dimension may be harder to isolate from other object-related dimensions. That is, in the real world, the position of a hammer within the periphery or the center of the visual field (i.e., its eccentricity) can easily be separated from its location in relation to the vertical meridian (i.e., polar angle) or its orientation. However, the typical grasp of a hammer (a power grasp) may be decodable also, to some extent, from the overall size of the object, potentially the amount of force needed to use the target object (e.g., precision grips may not be suited for high force grasps), or the weight of the object, among other mid-level (and potentially low-level) object properties. Nevertheless, and despite the fact that these dimensions are intertwined^1,14^, we can still disentangle them – e.g., the size of an object and its typical grasp are different types of information that can be independently impaired in individuals with brain damage^22,23^. This leads to the question of whether the organizational principles of sensorimotor cortices are also applicable to object-related information within association cortex. That is, are there true topographic maps for object-related dimensions – by analogy, contentopic maps – as there are for sensory dimensions (e.g., the eccentricity of a visual presented stimulus)?

## Modeling of object-related dimensions

To test whether object-related dimensions are represented topographically in the same fashion as low-level visual dimensions are, we presented participants with manipulable object stimuli in sequences that followed the rank-order of those objects in their scores on a particular object-related dimension^1^ while acquiring functional magnetic resonance imaging (fMRI) data. Specifically, we selected object-related dimensions that we obtained previously for a set of manipulable objects based on the subjective similarity between those manipulable objects on how they are manipulated in order to be used (see Methods)^1^. We chose the first two dimensions out of a 5-dimensional multidimensional scaling solution whose estimated distances between the tested objects were a good fit with the real distances (see Methods)^1^. That is, while these dimensions are still noisy and what they ultimately entail needs to be fully worked out, they are a relatively faithful description of the representational space we hold about how to manipulate objects. According to the labels provided by naïve participants, one of the selected dimensions refers to the type of grasp used when interacting with an object: power vs. precision grip (henceforth M1)^1^; the other dimension refers to the amount of force that is necessary when interacting with an object: dexterity vs. force (henceforth M2)^1^. Because we selected only the first two dimensions, perceived proximity between objects in this putative 2D space (M1 and M2) should be understood in the context of the full 5D space. Moreover, the names describing these dimensions, are, themselves, best approximations and may not fully capture the true underlying dimension, as they suffer from the noise inherent to the final multidimensional solution described above, as well as the necessarily subjective experience of the participants that labelled them. Nevertheless, at the level of analysis we are focusing here, these approximations are sufficiently detailed and have been shown to track, behaviorally and neurally, relevant object processing^1^.

Importantly, different objects have different scores for each dimension, and these scores refer to the “amount” each object has in relation to each dimension^1^. Thus, objects can be rank-ordered in terms of their scores per dimension. We took advantage of this, and in an experimental run (e.g., referring to dimension M1), we cycled 6 times through the object stimuli rank-ordered by the target dimension (e.g., from objects that are grasped with a clear power grasp to objects that are grasped with a clear precision grip; see Figure 1A & 1B, Figure S1, and Methods for information about the distribution of stimuli through a cycle as a function of the dimension being tested). What we are cycling through, then, are the different levels of the object-related target dimension, via the presentation of different objects according to the rank-order of their dimensional scores. What we are modeling is how cycling through these different dimensional levels elicits ordered changes in preferred responses in contiguous parts of the cortical surface in a continuous way. Objects were presented continuously and at the center of the screen. Importantly, M1 and M2 are mathematically orthogonal^1^, and thus object sequences for M1 are unrelated to those for M2. Moreover, we controlled for low level visual properties of the selected stimuli by covarying out potential contributions of eccentricity and polar angle of the different stimuli (calculated based on center of fixation and center of mass of each stimulus; see Methods) from our data, and used the residuals for our analysis. This is especially important because stimulus eccentricity is a potential contributor to the organization of mid-to-high level organization of object knowledge^16,19,21^.

**Figure 1.**
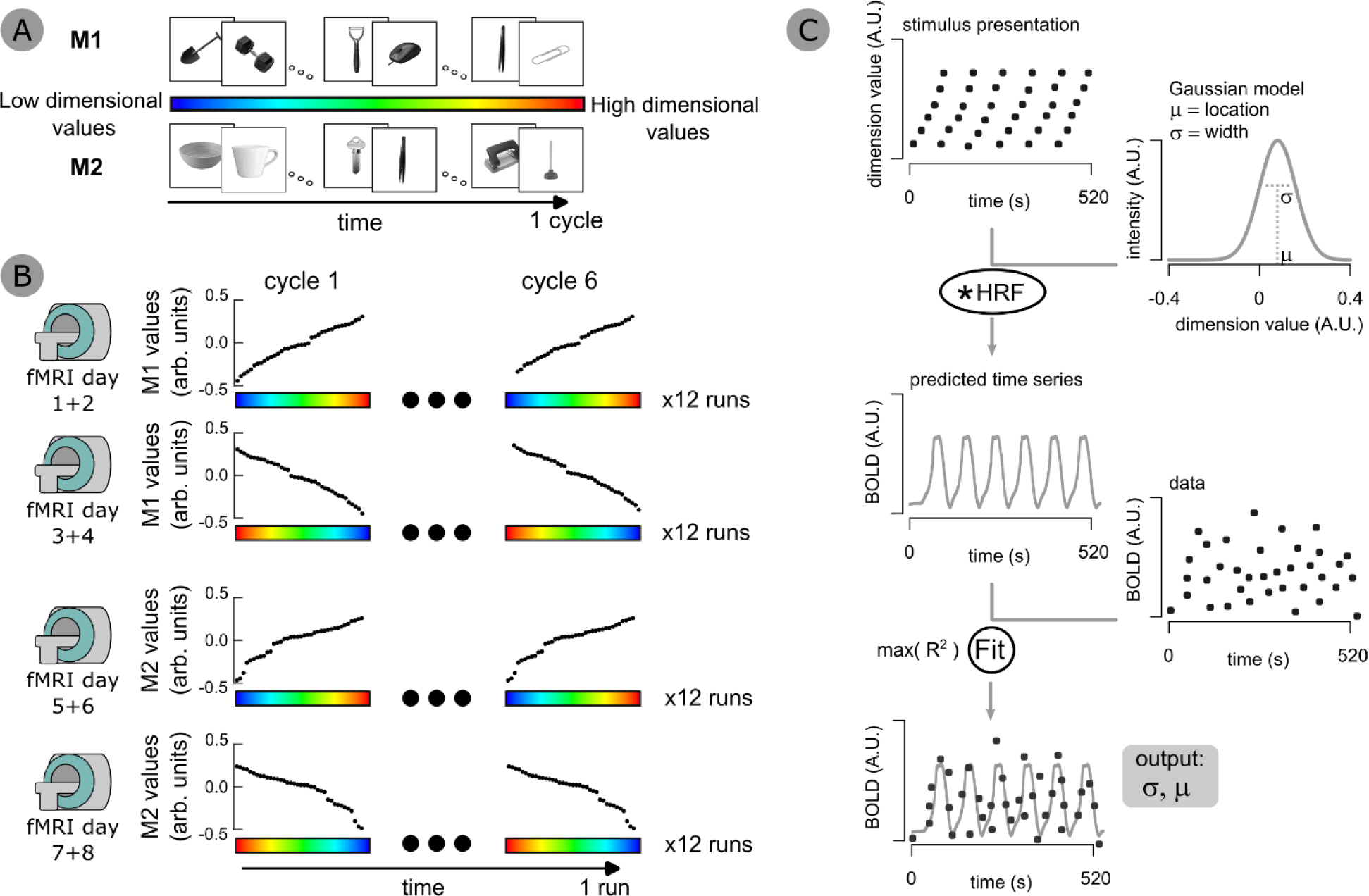
Experimental design and population receptive field modeling. Here we show (A) the progression of a cycle through different object types for each of the dimensions (M1 and M2). Forty stimuli were presented per cycle, sampling the different levels of the dimension evenly; (B) the distribution of the different cycles and orders per run and sessions. Participants went through 24 runs per dimension (in a total of 48 runs), 12 cycling the target dimension in one order (i.e., following ascending dimensional values), and 12 cycling in the reverse order. Each run cycled the dimension 6 times; (C) We adapted population receptive field modeling (pRF)^7^ to object-related dimensions. In this flow chart, we describe an example neural gaussian model for a particular voxel with particular tuning properties (preferred location in the dimension – µ – and tuning width – σ). The predicted time series for the selected voxel was calculated by passing the dimension-related stimulus presentation through the gaussian model. The resulting time course was then convolved with the hemodynamic response function. This created different fMRI predicted time courses for the different neural models tested, per voxel. Final model parameters – µ and σ – are obtained by minimizing the difference between the predicted and the observed fMRI data.

Given the two dimensions selected, potential contentopic maps should be obtained primarily within posterior parietal and dorsal occipital cortical regions, from the intraparietal sulcus posteriorly to V3d/V3A, as these areas code for shape and 3D representations of objects and grasp planning^24–28^. Moreover, aspects of object grasping that relate to typical object use (i.e., functional grasps)^29–31^ and to coding of grip force and object weight that allows for the derivation of the amount of force to apply to an object^32,33^, have been shown to be associated with responses within lateral and ventral occipitotemporal cortex starting posteriorly within lingual gyrus and fusiform gyrus.

To capture the systematic variation of fMRI signal elicited by the sequence of objects presented, as a proxy to the varying dimensional scores, we adapted standard population receptive field analysis (pRF) typically used to unravel the organization of low-level visual information (e.g., eccentricity)^7^, and applied it to our manipulable object dimensions. Specifically, we created one-dimensional gaussian pRFs to model neural signal (Figure 1C). The model had two parameters: one that coded for the preferred location in the dimension (i.e., which values of the dimension are preferred by the neural population being inspected); and one that coded the tuning width of the response (i.e., the specificity and range of dimensional values the neural population responds to). That is, we modeled preferred level of the target dimension (i.e., preferred location) across the cortical surface using a 1D gaussian with different tuning widths. Importantly, we show that this model reliably captures signal fluctuations as a function of the dimensional scores of the objects that were presented (goodness-of-fit: R^2^), accounting for the time course of different voxels by different preferred dimensional tunings and width values (Figure 2A, voxels in insets 1 to 3).

**Figure 2.**
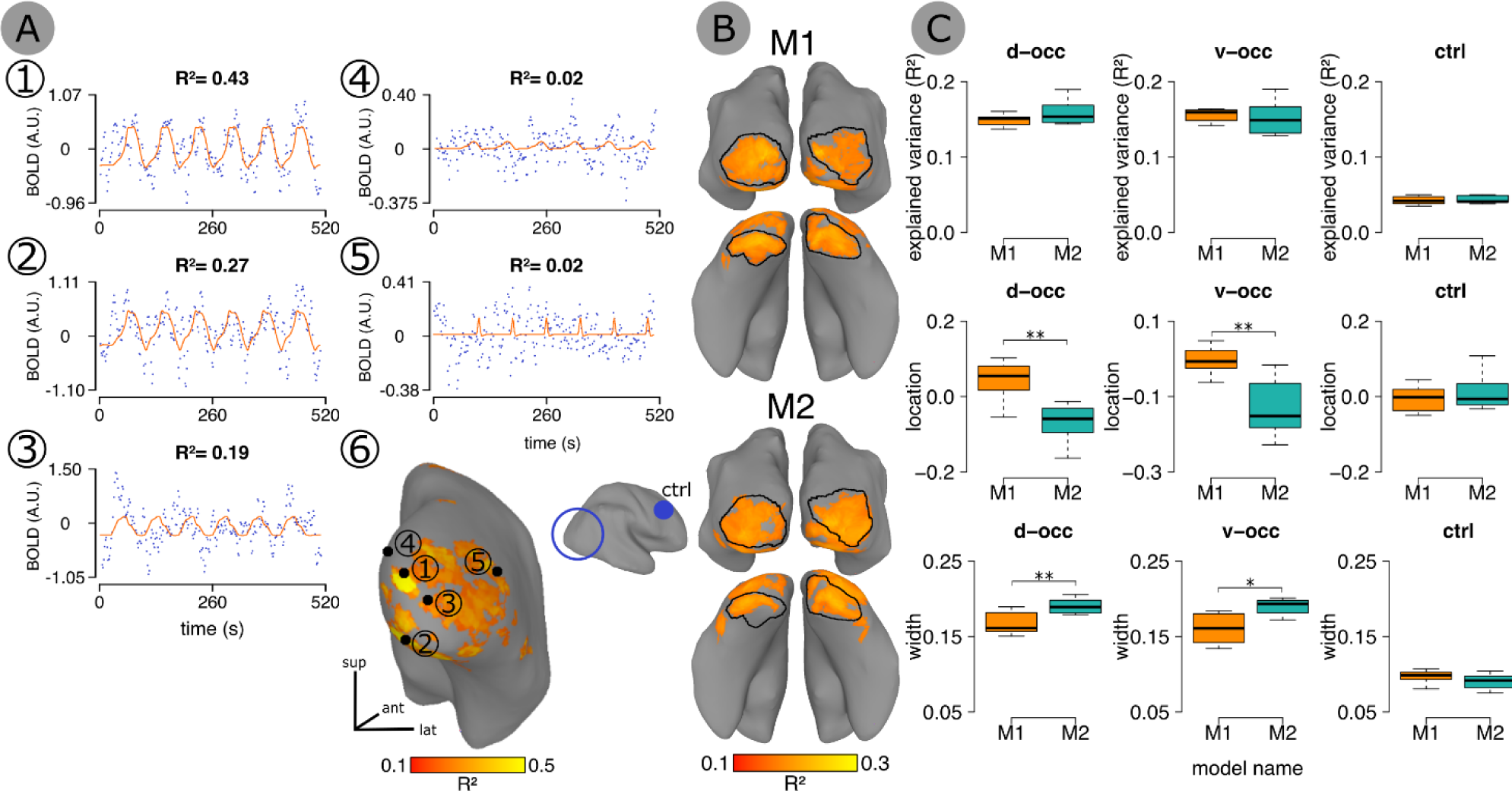
Population receptive field modeling of object dimensions, and neural population tuning. **(A)** Insets 1 to 5 show example fMRI time courses for different voxels for one dimension (M1). Inset 6 shows the location of the sampled voxels, as presented on the inflated right hemisphere of one participant. Each dot in the fMRI time course plot corresponds to blood oxygen level–dependent (BOLD) percent signal change. The red line corresponds to the model with the best fit that was attributed to the voxel, with the corresponding variance explained. Insets 1 to 3 show voxels whose time courses are well explained by the neural models created, whereas voxels 4 to 5 show voxels whose time courses are poorly explained by the neural models tested. **(B)** Group averaged goodness-of-fit maps of M1 and M2 dimensions. We defined our individual ROIs based on the four main clusters that survived thresholding at R^2^ = 0.10. We used a leave-one-participant-out approach to define the ROIs individually per participant and hemisphere. These include dorsal (d-occ) and ventral (v-occ) occipital association cortex. For visualization purposes, we delineated the ROIs based on the averaged individual ROIs. Moreover, we also created a control region of similar shape and size in fronto-lateral cortex (see Methods for details). **(C)** Distribution of goodness-of-fit (R^2^) and tuning parameters (preferred location in the dimension and tuning width) across participants for each dimension and each ROI. M1 and M2 yielded similar goodness-of-fit estimates across ROIs, that were systematically better than the goodness-of-fit estimates at the control ROI (ctrl, please note that data from the control ROI was not thresholded, as no voxels would survive a threshold of R^2^ = 0.10). M1 yielded an average higher location parameter compared to M2 across ROIs. M1 tuning was significantly lower compared to M2, indicating narrower tuning for the former compared to the latter. Moreover, and in line with the fact that, in general, extremely sharp tuning widths in pRFs are associated with noise-dominated data^7^, we obtained sharp tuning width estimates and low goodness-of-fit in the control region, in the context of width estimates in d-occ and v-occ that were considerably broader. Bonferroni corrected \**p*-values below 0.05; \*\**p*-values below 0.01; \*\*\**p*-values below 0.001.

We then tested how our model fares in relation to other potential alternative models. Specifically, we tested a purely linear model that predicts monotonical increases/decreases of neural response as a function of the dimensional scores (i.e., a linear model), as well as a simpler gaussian model that exclusively modeled preferred location in the dimension, using a fixed narrow tuning width (σ=0.05; effectively resulting in a location-only model). Our original 1D gaussian model with 2 parameters (i.e., preferred location in the dimension and tuning width) outperformed the other models in explaining the data (see Figure S2). As such, we focused exclusively on this model in all subsequent analysis exploring topographical representations of object knowledge.

## Comparing the tuning of the neural responses to the target object-related dimensions

In order to test for topography, we selected participant-specific cortical regions where our model robustly accounted for neural responses (R^2^ > 10%, approximately corresponding to p<10^-7^, uncorrected). This led to the definition of two bilateral regions of interest (ROIs): a dorsal occipital area (d-occ) around the dorsal aspects of the middle occipital sulcus and gyrus, including voxels in the vicinity of V3A^28^; and a ventral area around lingual gyrus, inferior occipital gyrus, and occipitotemporal cortex (v-occ; Figure 2B, Tables S1 and S2; Figure S3). Thus, the individual ROIs are generally within the regions expected to be tuned to these dimensions^22,24–33^. However, other cortical locations were engaged by the continuous presentation of objects in the particular orders dictated by different dimensions. Specifically, variations in the scores of M2 were also accounted for in bilateral regions in lateral temporal cortex (MNI coordinates for the left ROI −40, −70, −7, and for the right ROI 43, −69, −15). Finally, inspection of individual maps for both M1 and M2 demonstrates that most participants (6 out of 8; Figures S4 & S5) also showed responsive regions in posterior parietal cortex, but these cortical regions did not survive the strict inclusion criteria used.

We then focused on further understanding the topographical organization of the object-related dimensions of interest. M1 and M2 showed similar average goodness-of-fit of the pRF models across participants in d-occ and v-occ (Figure 2C, top inset), and these average goodness-of-fit values were much higher than those observed in a control region (Figure 2C, top inset; see Methods for the selection of the control region). Centrally, M1 and M2 differed in terms of average preferred location and width (i.e., the parameters of our pRF models) even though they engaged similar neural populations. This contrasts with the lack of a difference in location and width estimates for each of the dimensions in the control region (Figure 2C, middle and bottom insets).

Importantly, differences in average preferred location for each dimension at the two ROIs could not be explained by an overlap in the objects whose scores are within these average preferred locations for each dimension – the amount of object overlap around the average location estimated for M1 and M2 does not differ from what would be obtained by chance (permutation testing; see Methods; see Figure S6). Moreover, we observed a reliable difference in average width estimates between M1 and M2 within our ROIs. Specifically, M1 shows more specificity – i.e., its tuning is sharper – than M2. These patterns of results speak in favor of the reliability and sensitivity of our estimates.

## Object-related dimensions are arranged topographically

In the next step, we investigated the presence of contentopic maps – i.e., topographic maps that show a gradual and continuous progression of preferred dimensional scores over the cortical surface. We rendered preferred location maps for each dimension over the individual cortical surface (see Figure 3A for an example of an individual contentopic map for M1). To estimate the topographic organization of the maps, and to define their main axis, we conducted the following analysis: for each participant, each ROI, and each hemisphere, we computed a cortical distance map over the ROI along its longest axis, where 0 represents the middle of the ROI along this axis (Figure 3B left; see Methods for details). Each cortical distance map was rotated in 24 steps, yielding 24 cortical orientation maps (Figure 3B). We iteratively fitted each cortical orientation map with the preferred location map, for each dimension, to obtain the best linear relationship between the two. (Figure 3C). We then selected the orientations with the best fits per participant, dimension and ROI (Figure 3C, inset 2). Thus, these orientation maps capture the main axis of the contentopic maps.

**Figure 3.**
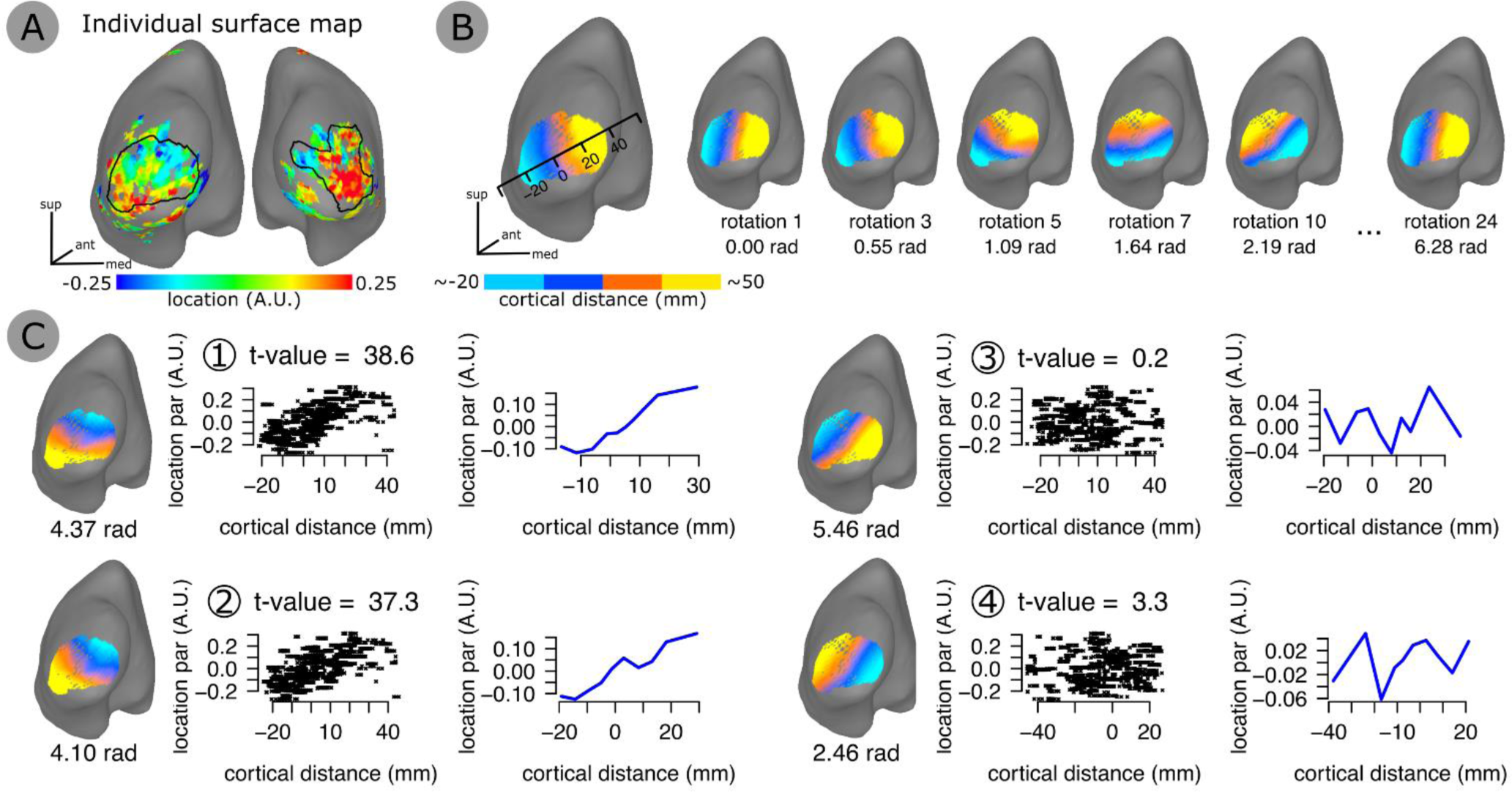
Estimation of topographical maps. **(A)** Gradual progression of the estimated location parameter on the d-occ ROIs for one participant, M1 dimension. **(B)** Cortical distance along the main ROI axis for one hemisphere (d-occ ROI) and multiple rotations between 0 (the output of the spectral decomposition, see Methods) and 2π (6.28 radians). **(C)** Four examples of cortical distance rotations and their relationship with the estimated location parameter that are shown in panel A. Panel C1 shows the best linear fit (t-value = 38.6) compared to panels C2, C3 and C4. Blue lines show the same data, but with the dimensional scores binned and averaged along 10 equally sized bins and shown across the cortical distance.

Figure 4A shows the average topographic progression and the average best orientation of the linear fits for dimensions M1 and M2. In both dorsal and ventral ROIs, we observed reliable topographical progression of preferred location within the target dimensions across the cortical surface – i.e., we obtained contentopic maps (see Figure S4 & S5 for individual best orientations of the linear fit). To quantify contentopic organization, we first showed that the best fits across participants are significantly stronger in d-occ and v-occ for both M1 and M2, when compared to those in the control region where overall poor goodness-of-fit was observed (Figure 4B). This demonstrates that there is a linear progression along the cortical surface as a function of the score of the objects in each of the dimensions.

**Figure 4.**
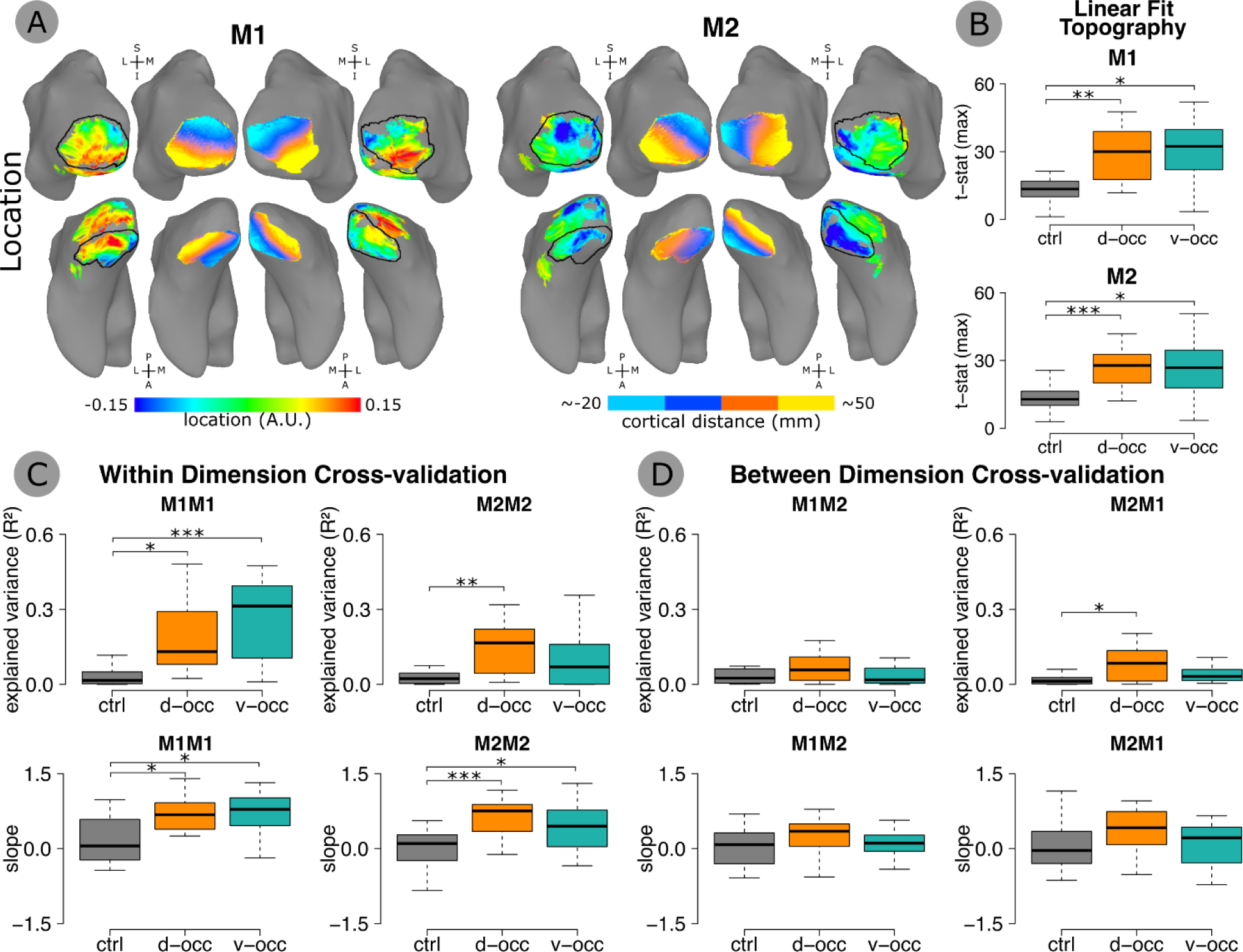
Contentopic mapping for location. **(A)** Average location parameter across 8 participants for M1 and M2 and average best orientations capturing the gradual representation of the location parameter. **(B)** Comparison of the t-statistic of the best orientation for each ROI (d-occ, v-occ and ctrl; please note that data from each ROI was thresholded based on the median R^2^ to provide a similar proportion of voxels across ROIs. Median R^2^ for d-occ and v-occ ∼ 0.11, median R^2^ for ctrl ∼ 0.04; see Table S3) and each dimension. Results indicate a better linear fit between cortical distance and location parameter in d-occ and v-occ for M1 and M2, compared to the control ROI. **(C)** Within-dimension cross-validation of the best location maps using a leave-one-participant-out procedure (see Methods). We assessed the similarity between the averaged best location map of the N-1 participants, and the best location map of the single-out participant using a linear regression model. We extracted the model explained variance (R^2^) and slope (beta coefficient). Note that a negative slope indicates estimates in opposite topographical direction (i.e., similar orientation but different direction of the map), whereas a positive slope indicates estimates in the same direction. Results indicate that we can successfully predict the single-subject best location map from the average or the N-1 group, for each ROI (d-occ and v-occ) compared to the control (ctrl). **(D)** Between-dimension cross validation. Same procedure as in C, but applied between dimensions. Results indicate that we cannot predict the best location map of the singled-out participant for one of the dimensions (e.g., M1) from the average of the N-1 group when using the best location maps of the other dimension (e.g., M2). This is evidence for the specificity of M1 and M2 topography, which is further corroborated by the difference in average location and widths parameter shown in Figure 2D.

## Object-related dimensions distinctively engage similar cortical locations

Finally, we tested the robustness of our fits for each dimension and each ROI via leave-one-participant-out cross-validation at the cortical surface level. If contentopic maps are consistent across participants, we should be able to predict the contentopic map of one of the participants from the contentopic maps of the remaining participants. Figure 4C (top insets) shows that the topographic progression of M1 and M2 location maps were consistent across participants, such that we can reliably predict the map on the left-out individual based on the average of the remaining participants. Importantly, we also tested cross-validation across dimensions – that is, we tested whether we can predict the contentopic map of one of the participants for one dimension (e.g., M1) from the contentopic maps of the remaining participants on the other dimension (e.g., M2). Consistency between participants was not found between the dimensions (Figure 4D). This shows, unequivocally, that these contentopic maps are dimension-specific – i.e., contentopic maps for M1 and M2 are distinct from one another.

However, the strongest test to the independence of the maps from M1 and M2 comes from analyzing the slope of the linear fits for the leave-one-participant-out cross-validation described above. This is because for the maps to be consistent across participants, they have to match in orientation, and most importantly, in the direction of the linear progression. In fact, simply matching in orientation could mean that the maps are in exact opposite directions. Thus, testing for positive slopes – i.e., when the to-be-predicted map and the maps of the remaining participants progress in the same direction through the cortical surface – will show that individual maps are consistent within dimension and independent across dimensions. The average slope of the within-dimension cross-validation linear fits within the ROIs for M1 and for M2 is positive (Figure 4C, bottom insets) and it is clearly different from the slope in the control region. This shows that, within each dimension, individual maps can be predicted from the maps of the remaining individuals both in orientation and direction of its topographical progression. This is in stark contrast with the between-dimension analysis. Here, slopes hover around 0 and are no longer different from the slopes in the control region. This demonstrates the reliability of the maps obtained for each dimension at the individual participant level, and, importantly, the independence of the contentopic maps for the two different dimensions (for the results on both explained variance and slope for the analysis of width of the tuning curve, please see Figure S7).

## Discussion

Previous studies have consistently demonstrated topographical mapping within sensorimotor cortices over particular dimensions that mirror response properties of the corresponding sensory receptor (e.g., location in the retina; sound frequency in the cochlea). This topographical organization is a central feature of sensory processing and dictates how we perceive and interact with the world^34^. Here we show that topographical organization is also present for more abstract kinds of information such as those pertaining to objects. Particularly, we obtained contentopic maps over object-related dimensions (e.g., grasp type) – i.e., neural responses within dorsal and ventral occipital cortex are tuned, independently, to the two object-related dimensions tested. These topographic maps are continuous, in that they progress through the cortical surface gradually as function of the level of the target dimension; are consistent across participants, in that maps of individual participants are predictable from the maps of the remaining individuals; and are specific for each dimension, in that topographic maps for one of the dimensions are not predictable from the maps of the other dimension.

The ROIs that show contentopy for the two target dimensions are part of visual association cortex. Importantly, while the content of the dimensions used herein still needs to be fully fleshed out, and the labels used are but approximations based on subjective judgements of naïve participants, generally, these dimensions describe object manipulation space. In line with this, the ROIs obtained have also been shown to code for action-related information. For instance, voxels within the dorsal occipital ROI are involved in the processing of object-specific 3D shape^27,28^, and participate in object grasping, and in hand shaping and orientation in preparation for grasping^22,26^. Moreover, voxels within the ventral occipital ROI have been shown to be also involved in object-specific (and potentially function-specific) action related processing^29,30^, as well as in the processing of object properties that impact action such as an object’s weight^32,33^. Undoubtedly, one should expect parietal cortical regions more anterior to the ROIs herein to also show some degree of tuning to the object-related mid-level dimensions used, as these areas code for object grasping and manipulation^25,26,35–38^. In fact, inspection of individual maps (Figure S3 & S4) shows that more posterior and superior parietal cortical regions do present neural responses tuned to both dimensions for most of the participants. Perhaps the weaker and less consistent responses in posterior and superior parietal cortex may be related with the format of stimuli presentation (exclusively visual) and the task required (unrelated to the object).

A remarkable aspect of our data is how independent the maps obtained for the two tested dimensions are. The way in which we elicited contentopic mapping reverted to using the same stimuli, with the only difference being that we presented these stimuli in different orders as dictated by the scores of each object in each dimension. Moreover, mapping was tested over the same neural population. Notwithstanding, pRF modeling of the neural responses (i.e., the modeling parameters of location and width) varied significantly (and non-accidently) by dimension. The computational separability of the neural tuning for these two dimensions suggests some level of topographic flexibility in how cortical regions process object-related information. Neural populations are able to pick up different informational properties of an object and code those independently in a superimposed way.

A major question relates to the nature of the representations within these contentopic maps. Our contentopic mapping data cannot be explained by low-level visual information. This is so because we regressed out stimuli-specific information about eccentricity (average pixel distance to center of fixation) and polar angle (orientation of the axis going from fixation to the center of mass of the object) from our data prior to the analysis reported here. Thus, contentopic maps were obtained independently of low-level stimulus variations of the kind that drive early visual cortex. This is not to say that low-level visual features, and especially eccentricity, are not informative and important for the definition of these maps and the dimensions that drive them (as they are of the response profiles of more anterior areas^16,21^) – for instance, eccentricity may well correlate, in the real world, with an object’s grasp. What we are showing, however, is that whatever is represented within these contentopic maps is not completely reducible to low-level properties.

Consequently, contentopy probably depends on intermediate representations that bridge between early visual properties and complex object representations^39^. Although these mid-level representations are important for visual and visuomotor object processing^40–42^, their nature is hard to determine^39^. Potentially, they no longer focus on spatial coding, but rather on object feature mapping^40,43^ and object-specific parameter spaces^44^. These spaces and maps should be dependent on the aggregation of information from upstream simpler cells (e.g., edge orientation) into more global representations (e.g., surface orientation). This would be done via highly non-linear transformations^4,39^ that group lower-level information using gestalt principles applied to visual and visuomotor^10,45^ processing. Interestingly, and related to the dimensions tested, models for object grasp have been proposed that rely precisely on gestalt principles to determine the kind of grasps to be applied to a stimulus through the computation of visual surfaces from low-level visual properties and focusing on action as a composite of chunks that respects surface properties^45^.

Speculatively, transformations over the readouts of the (multiple and potentially correlated) contentopic maps of the kind shown here would lead to composite higher order maps^4^ (within more anterior areas). In fact, it may be the case that domain selectivity within ventral temporal cortex follows precisely from the members of a domain satisfying a particular set of dimensions (and their associated contentopic maps; e.g., grasp type, elongation, presence of metal^1^) that relate strongly with that domain (e.g., manipulable objects). Thus, these downstream composite higher-level maps would arise dynamically from sampling upstream contentopic maps that are important for different domains of stimuli. This seems to be in line with there being finer parameter spaces within domain-specific regions^46^, and with the role of connectivity in the definition of the conceptual representation in ventral temporal cortex^3,47^.

But how does this possibility fit with many of the coarse topographies that have been suggested? Topography at more higher-level areas should emerge from the non-linear aggregation of upstream contentopy mid-level maps into composite maps^4^. For instance, domain specificity (and potentially real object size, a topography^15^ that is highly correlated with a domain organization)^2,3^ would emerge from the fact that members of a domain would evoke a large set of important dimensions, and thus mid-level contentopic maps, that are typical of that domain. In these domain-specific (or size-dependent) regions, traversing single dimensions (e.g., our M1 dimension) would lead to weak or no selectivity^4^. However, evoking the right pattern of dimensions would lead to the specific response preferences observed. Similarly, because non-linear integration of contentopic maps into composite higher-level maps follows dynamically from the requirements of the different domains, then regions that do not show particular domain preferences (the so called “NoManLand” regions^17^) should show more unidimensional maps, potentially tuned towards highly informative mid-level properties such as curvature singularities^10^, as they do^17,18^. Finally, this hypothesis is also compatible with the putative role of eccentricity at these higher-level topographies^16,21^. Eccentricity maps would work as protomaps^16^ to guide the dynamical integration of contentopies. Moreover, because contentopy is built out of from lower-level representations, eccentricity would be inherently passed to these higher-level maps.

Interestingly, this multidimensional flexibility in the topographical organization of object information may be central for the kinds of processes that underlie object recognition. The multidimensional flexibility that we show could be the mechanism by which object representations are built in the service of object recognition – neural populations would be tuned to different mid-level dimensions and flexibly superimpose different topographic maps. Potentially, these maps impose an object-centric reference frame to visual processing, just as retinotopy imposes a spatial reference to cognition^34^. One potential consequence of such composite maps is that they allow for flexibility and discriminability when processing individual items: a particular hammer may have some properties that differ from a prototypical hammer, but because of the intersection of the readouts of multiple contentopy maps, in the end, a hammer will still be a hammer, even if differing from a prototypical hammer in some dimensions.

Importantly, here we show that topography is (at least) one of the central organization principles that the brain uses to represent and store information about objects. Note that although we show contentopy for one type of objects – i.e., manipulable objects – and a particular set of dimensions, we predict that this contentopic organization should be present for other types of stimuli (e.g., faces, animals, scenes; provided that we have structuring dimensions for the object categories to be tested) and for other object-related dimensions. For instance, mid-level dimensions such elongation^1^, or particular surface properties^48^, should also lead to relatively independent contentopic maps.

Most importantly, our results demonstrate the ubiquity and non-incidental nature of topographical organization of information in the brain. Irrespective of the kind of information – whether sensorimotor^6,8,9^, or more abstract such as numerosity^49^ or, as demonstrated here, object information – the brain will organize information in continuous topographical maps that follow particularly relevant dimensional solutions for the kinds of information being coded. The parameters that may change for different kinds of information relate to the multidimensional flexibility that is implemented, and how this flexibility is rendered as a composite neural response. The benefits of such an organizational principle go from reducing metabolically costly long-distance connectivity between neurons processing similar information, to more cognitively focused aspects relating to processing specialization and stimuli discrimination^11,12^. Overall, our results raise the question of whether all kinds of information are amenable to topographical mapping. Although our data cannot fully address this, it clearly brings a new layer of understanding to both the general question of how information is processed in the brain, as well as to the quest of understanding what underlies our formidable capacity to recognize objects.

## Methods

### Participants

8 individuals (5 women; average age: 24; age range: 19-36) from the community of the University of Coimbra participated in the experiment. All participants were right-handed, had normal or corrected-to-normal vision, and were naive as to the experimental manipulations. The experiment was approved by the ethics committee of the Faculty of Psychology and Educational Sciences of the University of Coimbra and followed all ethical guidelines. Moreover, participants provided written informed consent, and were compensated for their time by receiving either course credit on a major psychology course or financial compensation.

### Stimuli

We used a set of 80 common manipulable objects previously used to determine and explore object dimensionality and for which we have scores in object-related mid-level dimensions^1^ (see Figure S1 for one exemplar of each object used). These were selected to be representative of the different types of objects that humans use routinely. We selected 3 exemplars per object type in a total of 240 images. Images were 400-by-400 pixel squares and subtended approximately 10° of the visual angle.

### Object-related dimensions

We selected two mid-level object-related dimensions from those that were obtained and tested in behavioral and neural experiments previously in our lab^1^. To extract these dimensions, we analyzed how different objects relate to each other in a large conceptual space, and extracted the main axes that organize this conceptual multidimensional space. Specifically, we presented 60 participants with words referring to the target 80 manipulable objects and asked them to think about how similar these objects were in the manner in which we manipulated them. We used an object sorting task to derive dissimilarities between our set of objects – each participant was asked to sort all 80 objects into different piles such that objects in a pile were similar to each other, but different from objects in other piles, on the target knowledge type. This piling solutions were then entered in a participant-specific similarity matrix, and an averaged dissimilarity matrix was then created^1^. We then applied non-metric multidimensional scaling (MDS)^50^ to the dissimilarity matrices obtained in order to extract key object-related dimensions that structure our object representational space. For details on the selection of the dimensions, the similarity matrices, and the behavioral and neural testing please refer to Almeida and Colleagues^1^. The two dimensions selected were the first dimensions, out of a 5-dimensional solution, that structured similarity in how we manipulate objects, and are the dimensions that explain most variance in these similarity judgements. The Kruskal stress value of the 5-dimensional solution was relatively low (Stress = 0,084) suggesting that the estimated distances between the tested objects in this multidimensional solution were a good fit with the real distances. These dimensions are by definition mathematically orthogonal. Moreover, these dimensions were labeled by another set of naïve participants as referring to the types of grasps applicable to the objects (M1), and to the amount of force that is necessary to interact with an object (M2; for details on all these experimental procedures and analysis please see Almeida and Colleagues^1^).

### fMRI Experimental design

The fMRI experiment consisted of 8 different sessions on different days. On day 1 and 2, dimension M1 was presented in a particular order. On day 3 and 4, dimension M1 was presented in reversed order in relation to sessions 1 and 2. On day 5 and 6, M2 was presented in a particular order, and on day 7 and 8 M2 was presented in reversed order in relation to sessions 5 and 6. Each session included 6 runs. Per run we presented 6 cycles of a dimension. For the first cycle we randomly selected 2 objects from each of the 20 bins (see Figure S1). The remaining 2 objects from each bin were then used in the following cycle. This means that per cycle we presented a sample of 40 out the 80 objects used to determine the dimensions, and that all the levels of the dimensions were evenly sampled. Specifically, for instance in an experimental run referring to dimension M1, we cycled through the object stimuli rank-ordered by this dimension 6 times. In a cycle, we first presented objects (one at a time) that are clearly used with a power grasp (e.g., a shovel, followed by a weight, followed by a tennis racket, etc.; see Figure 1A and Figure S1). These objects were then followed by objects whose grasp still falls closer to a power grasp than a precision grip (e.g., a grater, followed by a wooden spoon, followed by a computer mouse), and these were followed by objects whose grasp is increasingly closer to a precision grip (e.g., a fork, followed by a spray bottle, followed by a stamp). This cycle finished with objects that are clearly used with a precision grip (e.g., clothespin, tweezers and nail clipper).

Object images were presented for the duration of 1 TR (2 seconds). All runs were preceded and followed by 16 seconds of fixation. We used 3 different exemplar images per object which were counterbalanced across runs (exemplar 1 for run 1 and 3, exemplar 2 for run 2 and 5, and exemplar 3 for run 3 and 6). All participants completed all sessions and runs. Participants performed a simple task: they were instructed to maintain fixation on a yellow dot in the center of the image and press a button with index or middle finger when the dot changed color to red or green. Fingers and colors were counterbalanced across runs and sessions. The dot changed color on 10% of the trials.

### MRI acquisition

Scanning was performed with a Siemens MAGNETOM Prisma-fit 3T MRI Scanner (Siemens Healthineers) with a 64-channel head coil at the University of Coimbra (Portugal; BIN - National Brain Imaging Network). Functional images were acquired with the following parameters: T2* weighted (single-shot/GRAPPA) echo-planar imaging pulse sequence, repetition time (TR) = 2000 ms, echo time (TE) = 30 ms, flip angle = 75°, 37 axial slices, acquisition matrix = 70 x 64 with field of view of 210×192 mm, and voxel size of 3 mm3. Structural T1-weighted images were obtained using a magnetization prepared rapid gradient echo (MPRAGE) sequence with the following parameters: TR = 2530 ms, TE = 3.5 ms, total acquisition time = 136 s, FA = 7°, acquisition matrix = 256 x 256, with field of view of 256 mm, and voxel size of 1 mm3.

### Preprocessing of MRI data

Processing of the anatomical T1-weighted MR images was performed in Freesurfer image analysis suite (version 6.0, http://surfer.nmr.mgh.harvard.edu/) using the recon-all pipeline, that included white and grey matter segmentation, skull removal, and cortical reconstruction. Pre-processing of functional MRI data was performed in AFNI^51,52^ and included the following steps: slice acquisition time correction, head motion correction, detrending and scaling. Following the initial pre-processing steps, the time-series of each dimension (M1 and M2) and order of presentation (ascending and descending dimensional values) were averaged across repetitions for each participant. The averaged functional image was co-registered to the anatomical scans, using the function 3dAllineate, using local Pearson correlation as the cost function.

### Detrending low-level visual features

We calculated eccentricity and polar angle of the stimuli used in our experiment (3 sets of 80 exemplars) using Matlab (MathWorks). The image stimuli were first binarized to segregate the stimuli from the background. Eccentricity was calculated as the average distance of all pixels from the fixation point (corresponding also to the center of the image). To calculate polar angle, we first defined the center of mass of each object (using the regionprop function), and extracted the angle of the vector connecting the fixation point and the center of mass. The extracted values of eccentricity and polar angle were averaged across the three exemplars of each object, and across the 4 objects included in each bin. These averaging procedures were performed separately for each dimension. Thus, we obtained eccentricity and polar angle values for each bin of each dimension. These values were used to detrend the data, using the 3dDetrend function in AFNI, and the residuals of this detrending were then used in the pRF modelling procedure. By detrending these two dimensions we ensure that our maps are independent of the two major dimensions that lead to maps in visual cortex.

### Population receptive field analysis (pRF) and topography

The pRF analysis is a forward modelling approach aimed at accounting for the observed BOLD time series in each individual voxel by estimating parameters at the neuronal population level^7,49,53–55^. pRF parameters estimates were obtained from percent BOLD time series by fitting a linear model via an iterative procedure, similar to previous applications in the domains of vision, sensorimotor integration and quantity/numerosity perception The linear model can be described as follows:

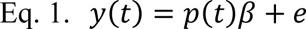

In equation 1, *p(t)* represents the predicted BOLD signal, β is a scaling factor necessary to match the prediction to the observed data, defined in arbitrary units (A.U.), and *e* is measurement error, representing the departure from the estimated and the observed BOLD. The fitting was performed via the General Linear Model approach (GLM), using ordinary-least-square estimation (OLS).

We obtained the prediction *p(t)* starting from a parameterized model of the stimulus sequence. To parametrize the stimuli sequence, we used a Gaussian tuning function (see equation 2) with the neuronal parameters (location, µ; width, σ) arranged in a two-dimensional grid. Gaussian location parameter (µ) was allowed to vary between −0.6 and 0.6 in 20 steps. Gaussian width (σ) was allowed to vary between 0.05 and 0.25 in 18 steps, for a total of 360 predictions.

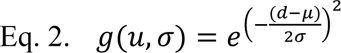

For each voxel’s percent BOLD time series, the fit was performed with an iterative procedure. On each iteration the predictor was built based on the selected width and location parameters. The predictor was set as the independent variable of a general linear model with the percent BOLD time series as the dependent variable. For each iteration we computed model goodness-of-fit, measured with R^2^. The set of neuronal parameters (location, µ; width, σ) that maximized R^2^ were assigned to the individual voxel.

It is important to note that in this modelling approach, the estimated parameters (Gaussian location - µ; and width - σ) are connected to local features of the percent BOLD time series. This approach facilitates the following analyses (statistical and topographical), allowing to assess the distribution of specific features of the percent BOLD time series across different experimental conditions and ROIs.

### Definition of ROIs

For each individual, we defined the regions of interest (ROIs) based on a leave-one-out procedure. For participant *x*, we projected pRF parameter maps (R^2^, location and width) on standard surface for each of the remaining (n-1) participants using the AFNI function 3dVol2Surf – i.e., we create 8 n-1 participant averages, one for each left-out participants thus defining independent ROIs.

Compared to standard volumetric approaches, the standard surface maps approach offers an advantage in terms of between participant co-registration at the cortical level^56^. Surface-based mapping relies on matching between common anatomical features such as sulci and gyri, and it preserves the local topology of the individual cortical sheet at a single hemisphere level. Thus, this procedure is more accurate than what could be obtained by classical volumetric-based registrations between individuals.

We averaged and visualized the maps obtained from the n-1 participants in the common MNI surface space. For each hemisphere and tested dimension (M1 and M2), we average R^2^ across the n-1 participants and threshold the resulting map above 10% R^2^. We then ran a clustering algorithm at the surface level to remove small clusters of activation that survive the thresholding (minimum cluster size: 35mm^2^). The outcome of this operation yielded two ROIs per hemisphere: a dorsal-occipital ROI (d-occ) and a ventral occipital ROI (v-occ). For visualization purposes we then averaged these ROIs. Please see Tables S1 and S2, as well as Figure S3 for the location of these averaged ROIs.

The d-occ and v-occ rois were then projected back on the left out individual x for subsequent analysis. Please note that all the analyses presented in the current manuscript were performed on the native and topologically correct individual participant space.

### Permutation testing

Within the ROIs, the best modelling parameters across participants differed between object-related dimensions M1 and M2 (Figure 2C, main manuscript). Specifically, M1 yielded more positive tuning location estimates and narrower tuning width compared to M2. From an fMRI perspective, the only difference between M1 and M2 is the temporal order in which the visual stimuli are presented. It might be argued that the observed differences between M1 and M2 modelling parameters could be simply due to a response to particular objects within the sequence, presented at different times during each dimension-specific sequence.

To test for this possibility, we extracted 12 objects around the average location estimate of M1 and M2 (i.e., 3 bins), separately. We observed an overlap of 2 out of 12 objects between M1 and M2 (16.6% of objects overlap). We asked whether this overlap would be sufficient to drive the observed difference between M1 and M2 average location estimates. To assess this, we run a permutation test, where we obtained the null distribution of the percentage of object overlap between 12 elements extracted separately from two shuffled sequences of 80 objects. If obtaining a 16.6% overlap would be extremely rare, then it could be argued that the difference between M1 and M2 average location estimates might be driven by object overlap per se. If 16.6% would fall within the expected range of overlap, then it would be less likely that the object overlap would account for the M1 and M2 location difference.

Permutation results indicate that the 95% confidence interval of the null distribution ranges between 0% and 33%, with a median of 16.6%. That is, in 95% of the repetitions, the number of overlapping objects between the two sequences ranged between 0% and 33% (see Figure S6). In our data, we observe an overlap of 16.6% between the objects around the average location parameter of M1 and M2 – which falls within the 95% confidence interval of the null distribution. Thus, the number of objects overlap derived from our data is not different from what we would expect by chance. Hence, the difference in tuning parameters we observe is unlikely to be accounted for by the number of overlapping objects falling around the estimated mean tuning locations.

### Analysis of the orientation of the contentopic maps

A topographic map is a gradual and ordered representation of an estimated variable over the human cortical or subcortical location (e.g., visual eccentricity^12^). To test for the presence of topographical maps at the individual participant level we proceeded as follows: for each participant’s ROIs and hemisphere we computed the node-to-node geodesic distance from each node in one ROI – the adjacency matrix, derived from the distance metric, *dist –* geodesic distance, between two mesh vertices *x_i_* and *x_j_*:

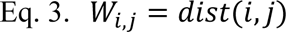

Then we computed the spectral decomposition of the adjacency matrix and selected the first eigenvector of the decomposition (vector 1 – *v_1_*)^57^. This vector represents a map of cortical distance over the ROI along its longest axis, where each point in the map is the geodesic distance of a node along the selected axis with respect to 0 – the middle of the ROI along its longest axis. The second eigenvector of the decomposition (vector 2 – *v_2_*) represents a map of cortical distance over the ROI along its shorter axis.

Vector 1 and vector 2 effectively represent a 2D mapping over the cortical surface. These are used for example in surface-flattening approaches. In this application we iteratively rotated the 2D mapping with respect to its original axis in 24 steps using an 2D matrix rotation of the desired angle (in radians), multiplied by the original 2D mapping coordinates (equation 4). Note that this operation does rotate the vectors in surface space but does not shift the center of the vectors.

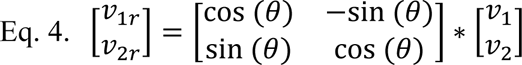

For each iteration, this operation resulted in a rotated version of vector 1 and vector 2 (resulting in rotated vectors – *v_1r_ v_2r_*). We used vector 1 as our reference for the subsequent analysis steps.

For each participant, ROI, dependent variable (pRF location and pRF width) and dimension tested (M1 and M2), each of the 24 rotations of vector 1 map were projected in the single participant anatomical space using the AFNI function 3dSurf2Vol. We compared the linear progression between the rotated vector 1 map and the estimated dependent variable. The comparison between the rotated vector 1 and the variable of interest was performed using simple linear regression. The obtained t-statistic was stored for subsequent analysis.

For each participant, ROI, dependent variable (pRF location and pRF width) and experimental condition (M1 and M2) the best orientation characterizing the map was defined as the one yielding the highest t-statistic among the 24 possible rotations (linear fit – topography, see main manuscript, Figure 4B). Following this procedure, we obtained the best rotations capturing the gradual representation of the given estimated variable (pRF location and pRF width) over each participant human cortical surface and ROI. We compared the obtained highest t-statistic for each participant and ROI with those obtained from a control ROI of similar size derived from each individual dorso-lateral cortex (ctrl ROI, main manuscript, Figure 2B and Figure 4B).

We opted for a linear model for theoretical (i) and computational (ii) reasons. (i) Theoretically, topographic maps are, for the most part, arranged linearly over the cortical surface, with a notable exception being tonotopic maps showing a non-linear progression along the Heschl’s gyrus (Da Costa et al., 2011). Somatotopic maps, although showing a discontinuity necessary to cover the two-dimensional body over a quasi-1D strip of cortex (pre-central and post-central gyrus), are largely arranged linearly when taking one effector at a time. For example, the hand shows an ordered progression of finger representation over the cortical surface^58^. Thus, at the very least, linear models would be best approximations for a first level analysis of contentopy; (ii) From a computational perspective, adding non-linear components would increase the likelihood of overfitting, thus potentially capturing noise variability instead of ‘true’ underlying map features.

### Cross validation of contentopic maps

We cross-validated the gradual representations for each ROI and each participant using a leave-one-participant-out procedure. For each iteration we averaged the map of N-1 participants (either location or width) in MNI surface space and compared the corresponding map (location or width) of the left-out participant. We assessed the similarity between the averaged map of N-1 participants and the left-out participant map using a linear regression model, storing the model goodness-of-fit (R^2^ – explained variance) and the model slope (beta coefficient). We report the model goodness-of-fit and model slope, as the linear model could yield high explained variance (R^2^) estimates even in if the averaged map of N-1 participants and the left-out participant map were going in the opposite direction. Reporting the model slope resolves this issue, as the sign of the slope indicates the direction of the relation between maps. A negative slope indicates estimates in opposite direction, a positive slope indicates estimates in the same direction.

## Acknowledgements

This research was supported by the European Research Council (ERC) under the European Union’s Horizon 2020 research and innovation programme Starting Grant 802553 “ContentMAP” to JA, and by European Research Executive Agency Widening programme under the European Union’s Horizon Europe Grant 101087584 “CogBooster” to JA. SK was supported by a Fundação para a Ciência e Tecnologia (FCT) Doctoral Grant SFRH/BD/145218/2019, ZT was supported by a Postodoctoral fellowship under the European Research Council (ERC) under the European Union’s Horizon 2020 research and innovation programme Starting Grant 802553 “ContentMAP”. AF was supported by a Biotechnology and Biology research council (BBSRC) grant BB/S006605/1, and a Bial Foundation Grant 203/2020.

## Author contributions

Conceptualization: JA Methodology: SK, AF, ZT, JA Investigation: SK, ZT Funding acquisition: JA Analysis: AF, SK, ZT, JA Writing – original draft: JA Writing – review & editing: AF, SK, ZT, JA

## Code Availability

Codes will be available at OSF before publication.

## Data availability

Data will be available at OSF before publication.

## Extended data and figures

**Figure S1.**
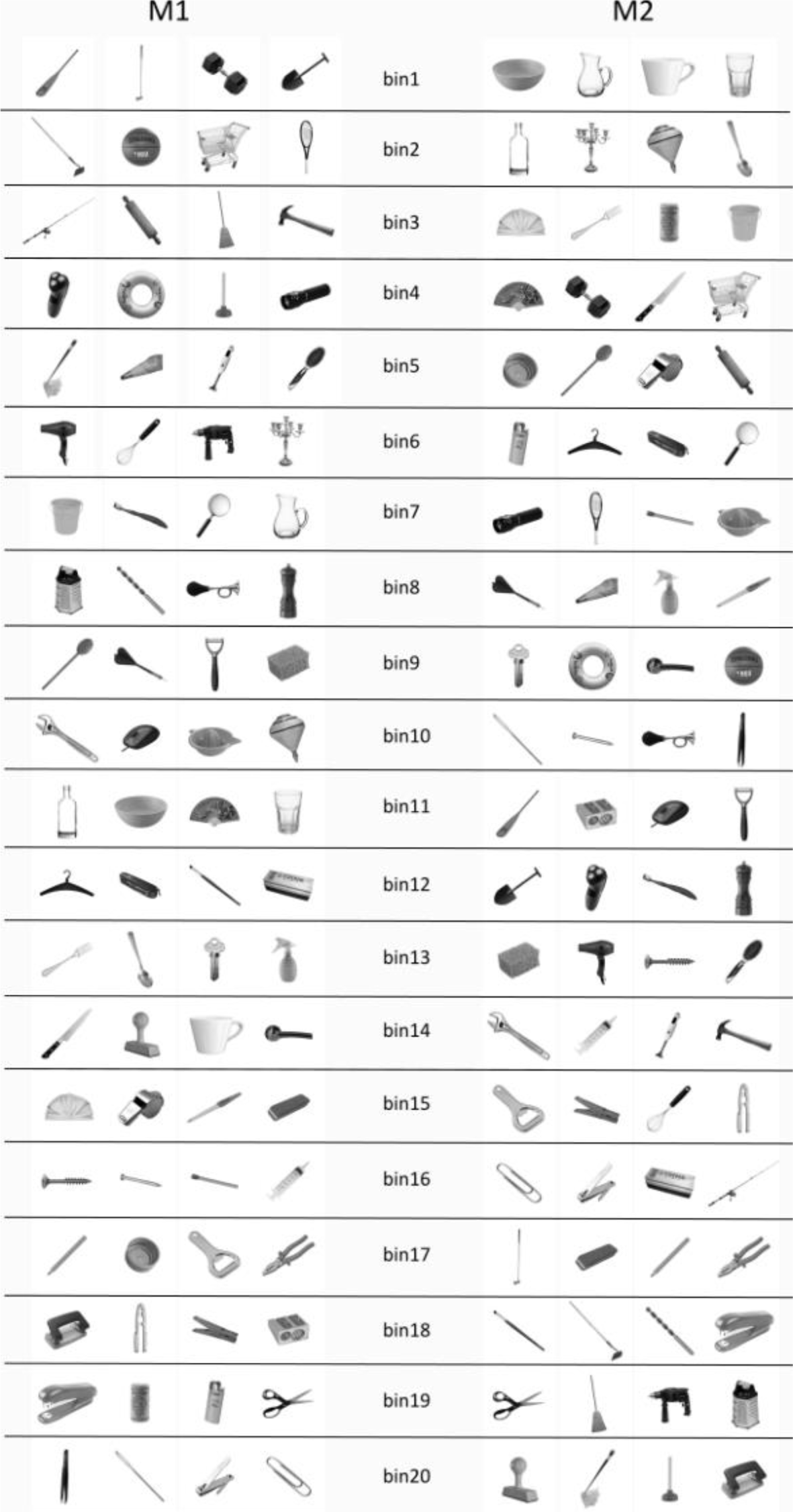
Examples of the stimuli used and their rank-order in the target dimensions. Here we present one exemplar image per object used, as well as the rank order of each object per M1 and M2. Each object has a score per dimension extracted. To help with the design and the cycling of each dimension through all its levels evenly, we grouped the objects in twenty bins. These bins were sampled evenly in each of the 6 cycles of a run.

**Figure S2.**
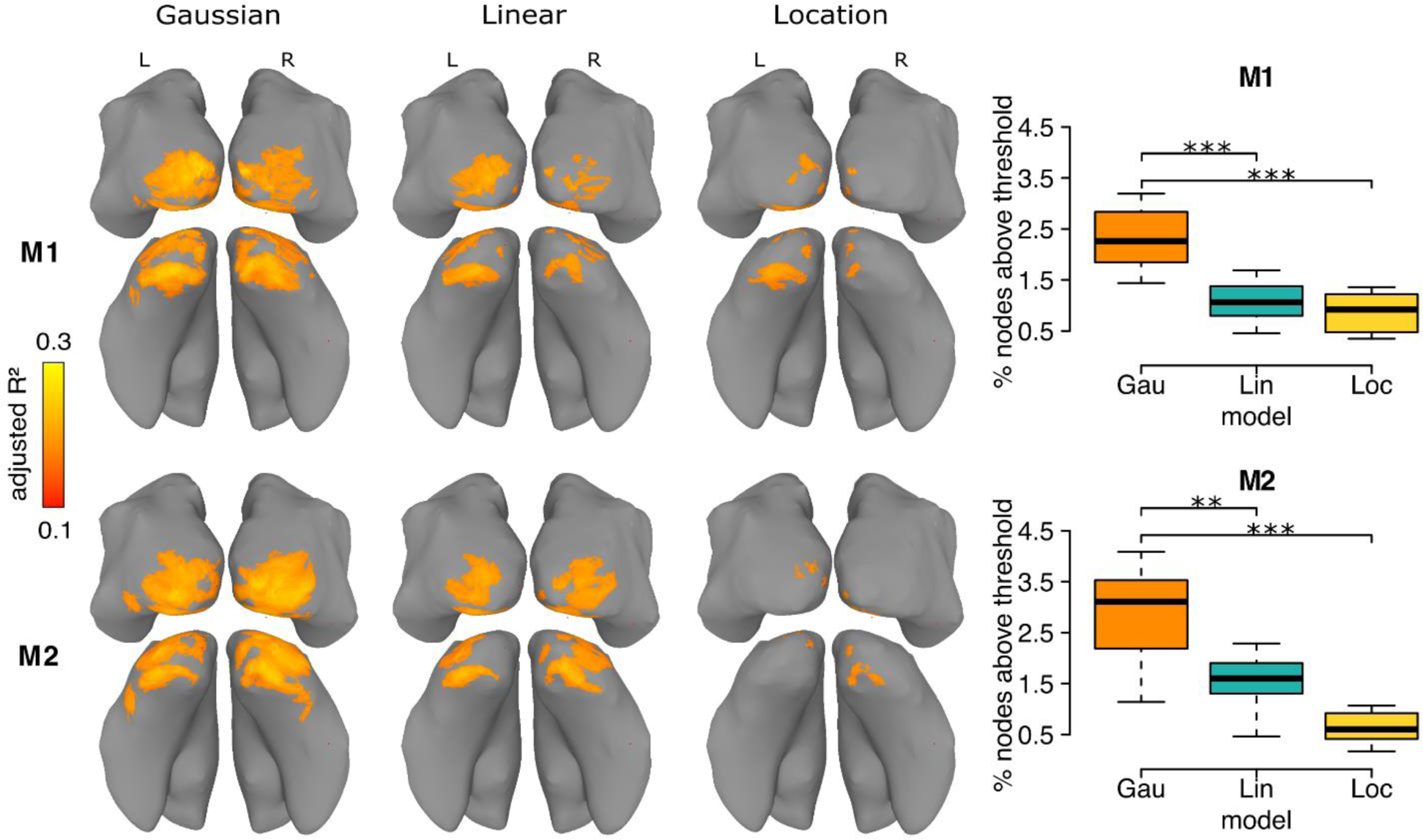
Comparison of the 1D gaussian model with two parameters with alternative models. We compared the goodness-of-fit of the 1D Gaussian model with two parameters with the goodness-of-fit of two alternative models. Here we show maps of the adjusted R^2^ values and the direct comparison in the percentage of nodes above the threshold. This clearly shows that the 1D Gaussian model with two parameters outperforms both the Linear and the Location only models.

**Figure S3.**
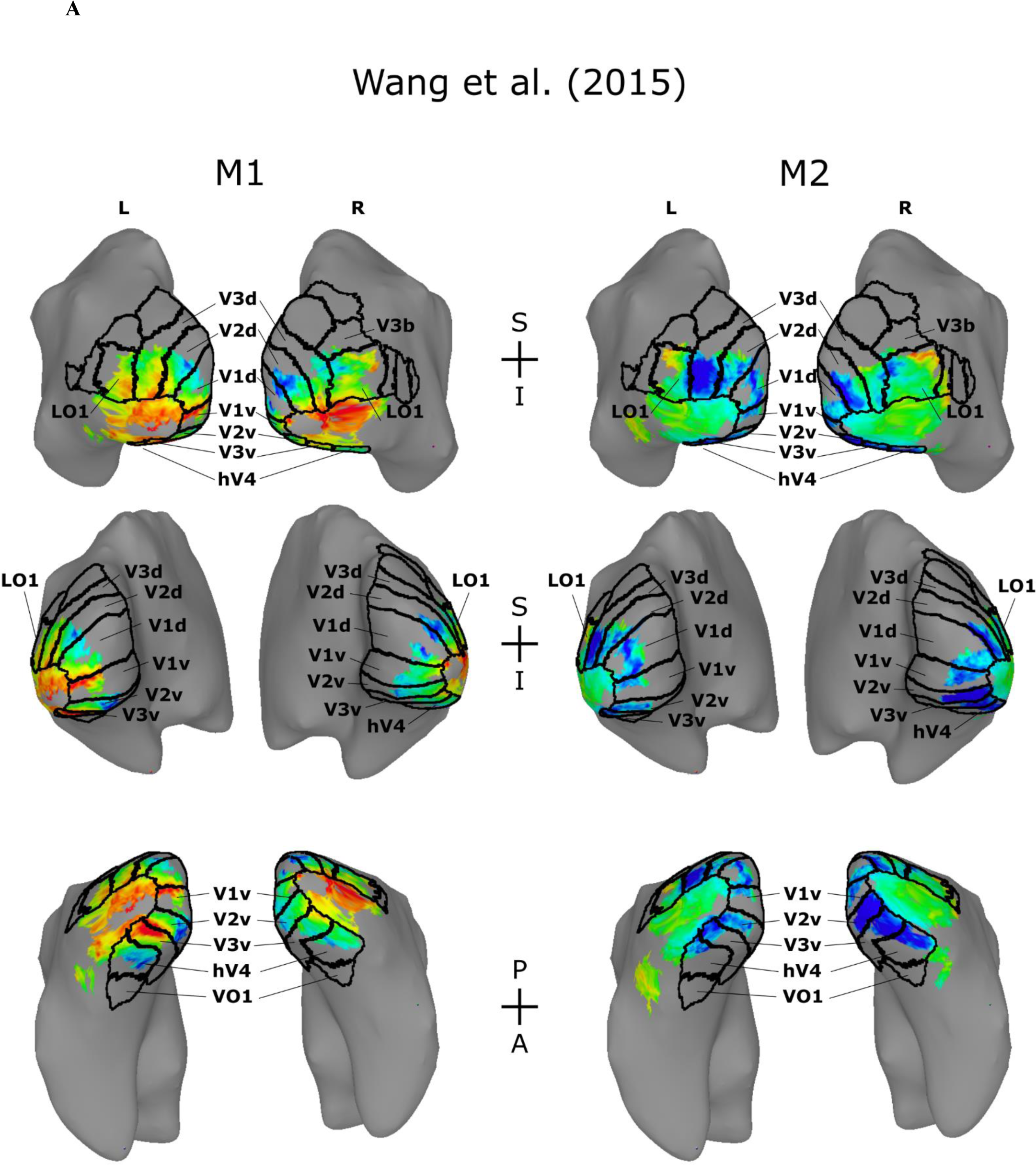
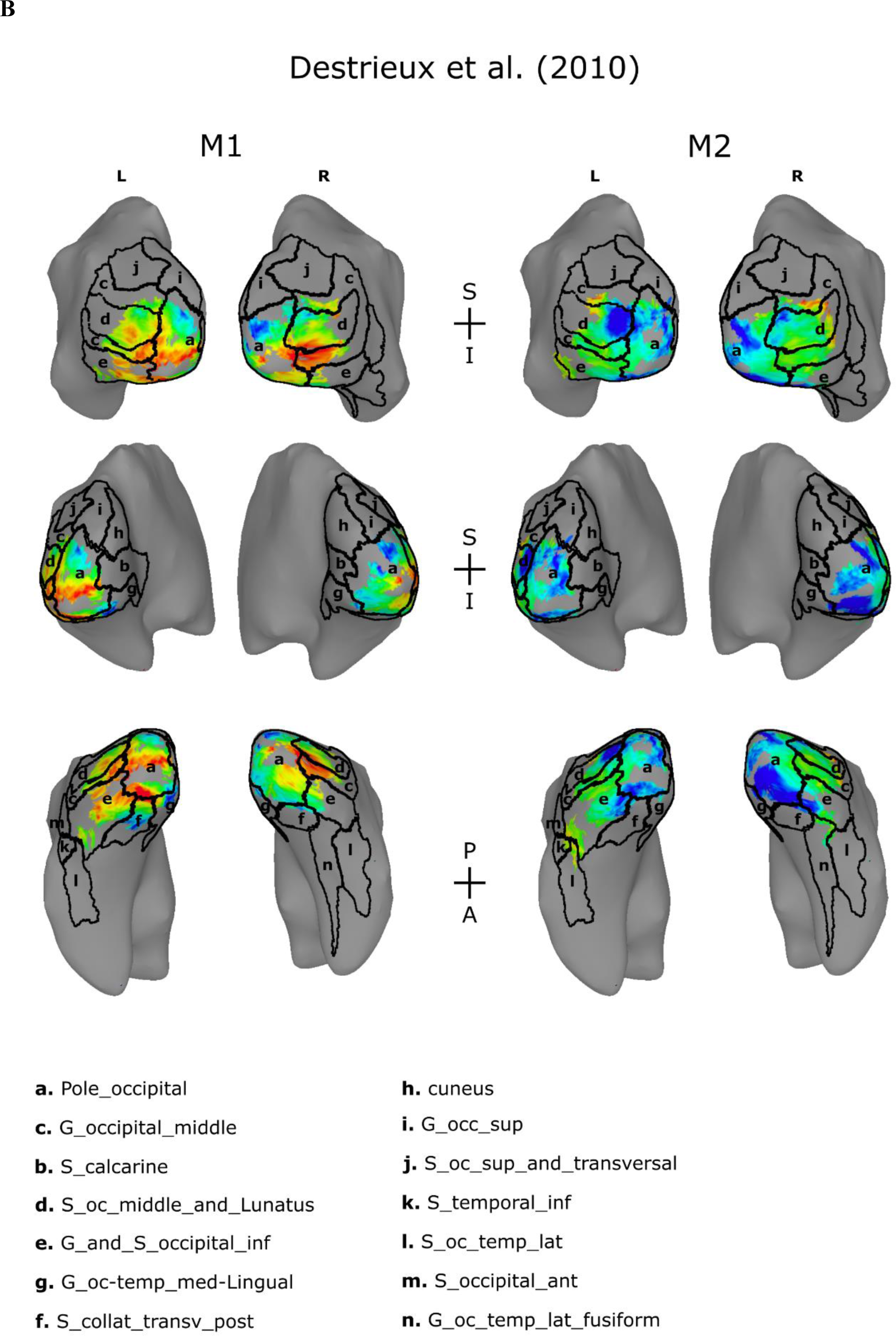
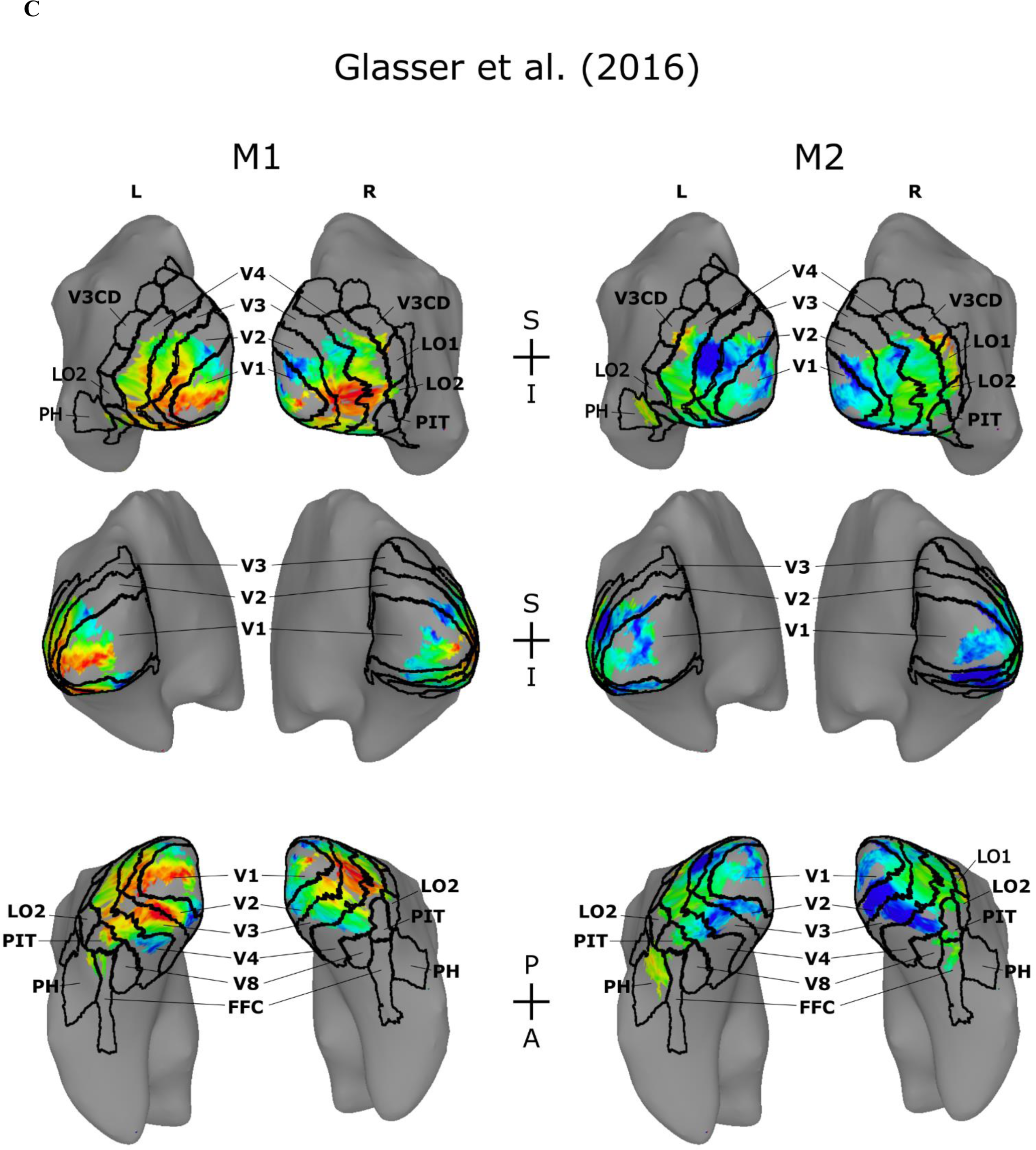
Average contentopic maps for M1 and M2 with atlases. Maps for the location parameter across the 8 participants with overlapping regions from 3 different atlases outlined in black. (A) Wang et al.^59^; (B) Destrieux et al.^60^; and (C) Glasser et al.^61^.

**Figure S4.**
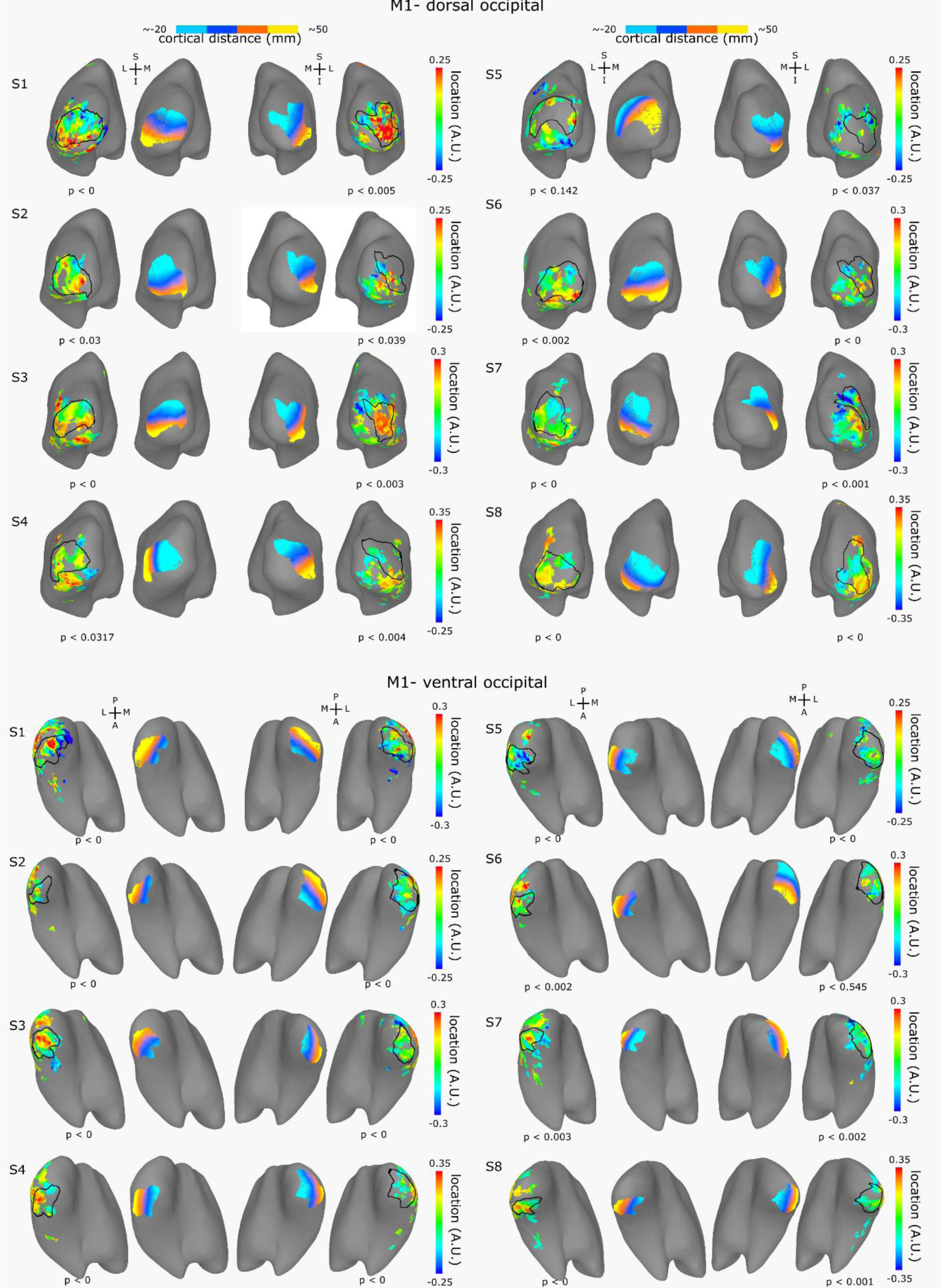
Individual Contentopic maps for orientation for M1. Maps for the location parameter across the 8 participants for M1 in dorsal and ventral views.

**Figure S5.**
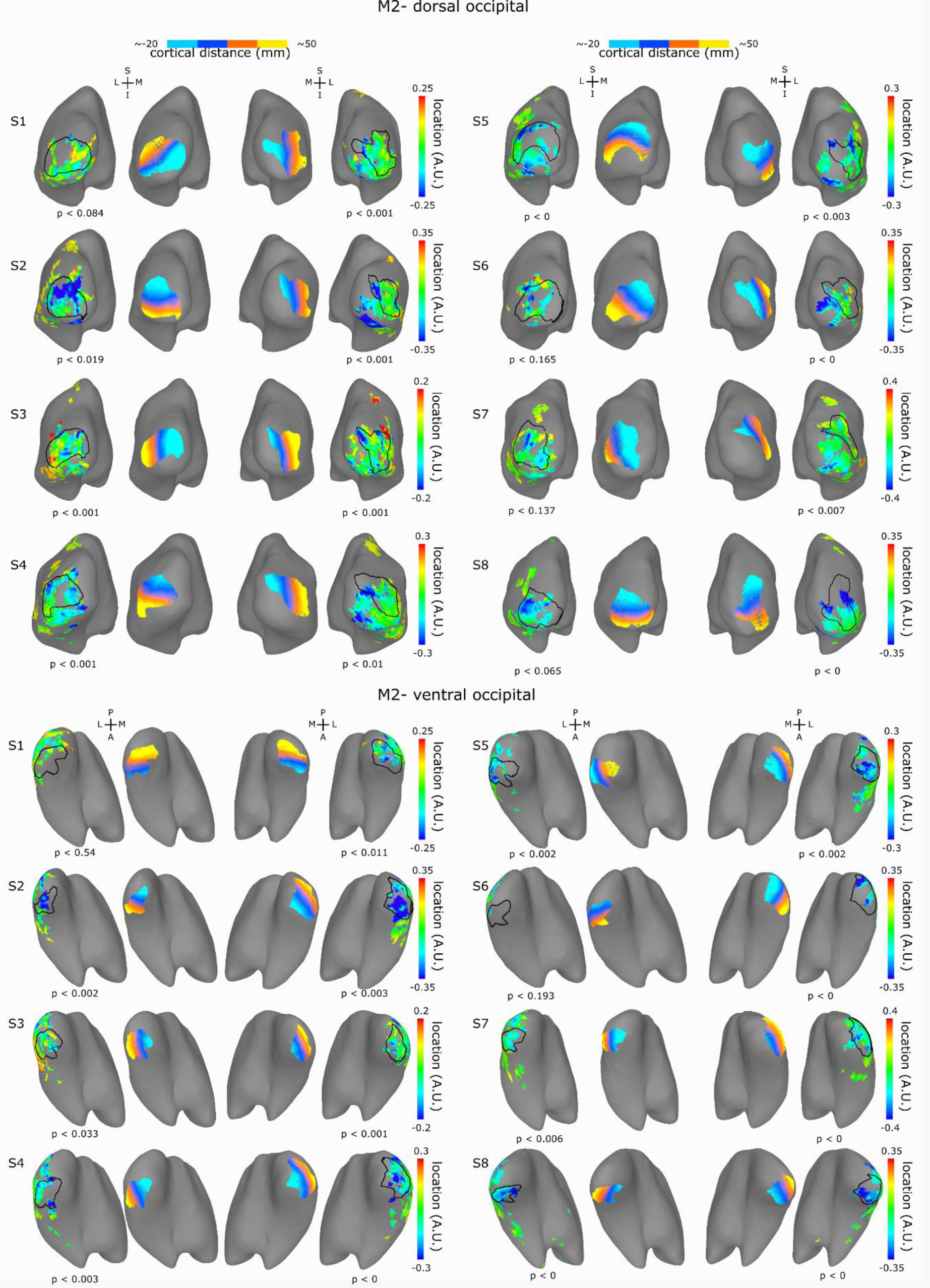
Individual Contentopic maps for orientation for M2. Maps for the location parameter across the 8 participants for M2 in dorsal and ventral views.

**Figure S6.**
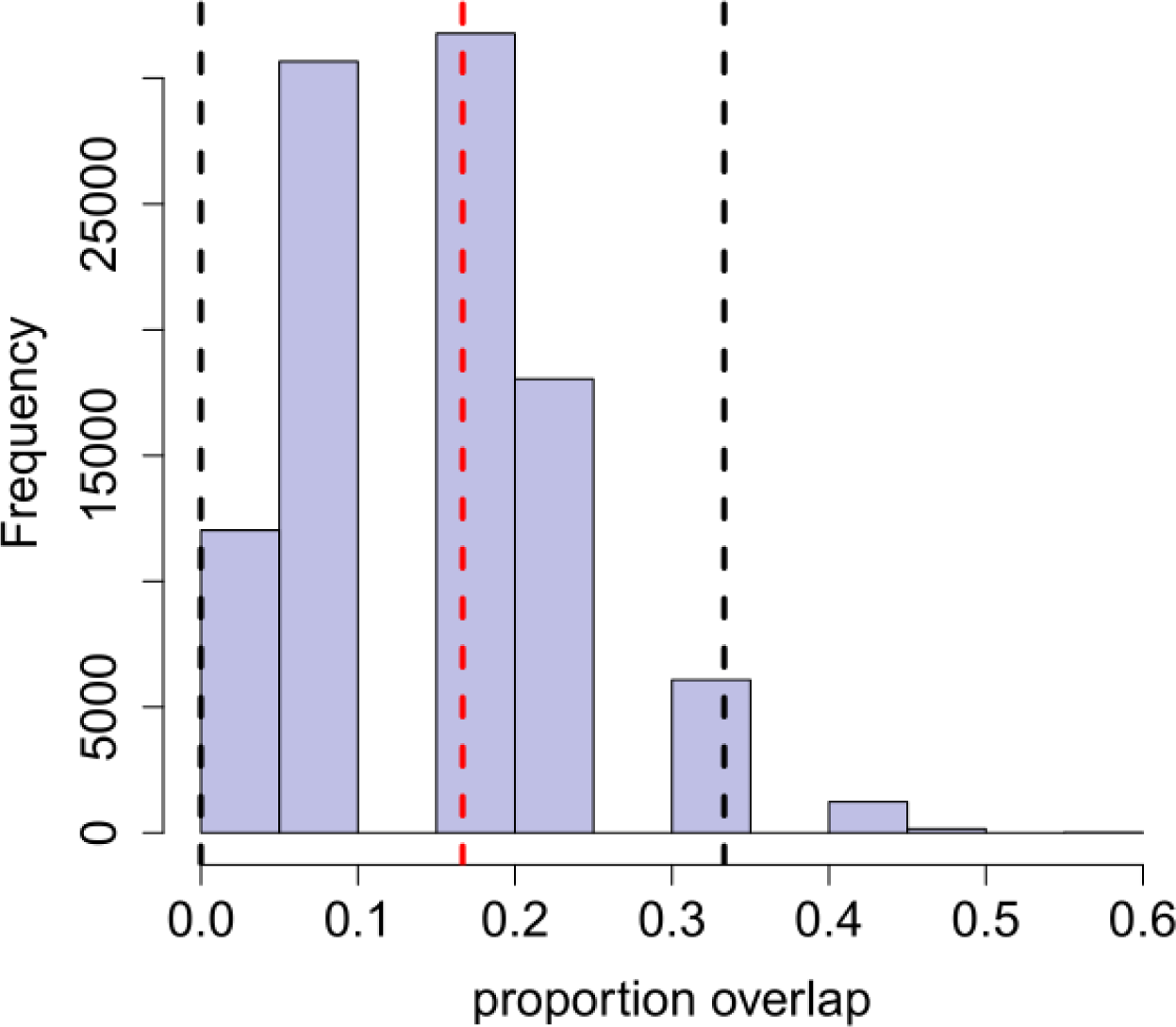
Distribution of the permutation test of object overlap within the average location values for M1 and M2. The figure presents the null distribution of the percentage of object overlap between 12 elements extracted separately from two shuffled sequences of 80 objects. The observed overlap found in our data (red dashed line) falls within the 95% confidence interval (black dashed lines), indicating that the overlap we found here is not different from chance level.

**Figure S7.**
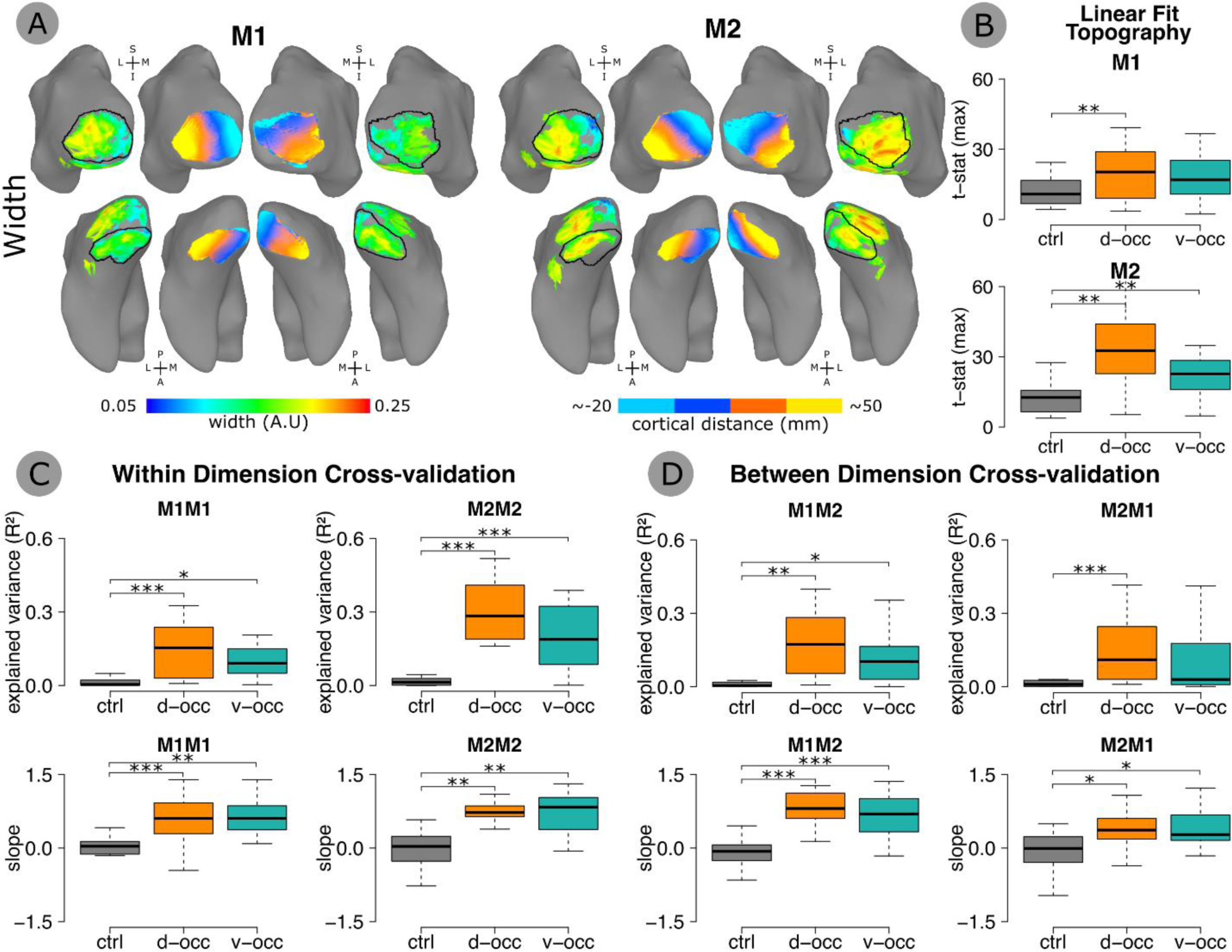
Contentopic mapping for width. (A) Average width parameter across 8 participants for M1 and M2 dimensions and average best rotations capturing the gradual representation of width parameter. (B) Comparison of the t-statistic of the best orientation for each ROI (d-occ, v-occ and ctrl) and each dimension. Results indicate a better linear fit between cortical distance and width parameter in d-occ for M1 and M2, and v-occ for M2 compared to the control ROI. (C) Within-dimension cross-validation of the width maps using a leave-out procedure (see Methods). We assessed the similarity between the averaged width maps of the N-1 procedure, and the width map of the singled-out participant using linear regression model and extracting the model explained variance (R^2^) and slope (beta coefficient). A negative slope indicates estimates in opposite direction, a positive slope indicates estimates in the same direction. Results indicate that we can successfully predict the singled-subject width map from the average or the n-1 group, for each ROI (d-occ and v-occ) compared to the control (ctrl). (D) Between-dimension cross validation. Same procedure as in C, but applied between dimensions. Results indicate that we can predict the single-subject width map from the average or the n-1 group when using M1 data to predict M2 or vice-versa.

**Table S1.** Atlas overlap. Regions of overlap between the average contentopic maps and the Wang et al.^59^, Destrieux et al.^60^, and Glasser et al.^61^ atlases. Region names are according to the atlas naming convention and are followed in brackets by their numerical code from the atlas. Region names from the Destrieux et al.^60^ atlas are preceded by “S_” and “G_” to indicate sulci and gyri, respectively.

| Wang |  |  |
| --- | --- | --- |
| Left | *d-occ* | V1v (1), V1d (2), V2d (4), V3d (6), LO1 (15), LO2 (14) |
|  | *v-occ* | V1v (1), V2v (3), V3v (5), hV4 (7) |
|  | *ctrl* | no overlap |
| Right | *d-occ* | V1d (2), V2d (4), V3d (6), LO1 (15) |
|  | *v-occ* | V1v (1), V1d (2), V2v (3), V3v (5), hV4 (7) |
|  | *ctrl* | no overlap |
| Destrieux |  |  |
| Left | *d-occ* | Pole_occipital, G_occipital_middle, S_oc_middle_and_Lunatus, G_and_S_occipital_inf |
|  | *v-occ* | Pole_occipital, G_oc-temp_med-Lingual, S_collat_transv_post, G_and_S_occipital_inf |
|  | *ctrl* | S_front_middle, G_front_middle, S_front_middle, S_front_sup |
| Right | *d-occ* | Pole_occipital, G_occipital_middle, S_oc_middle_and_Lunatus, G_and_S_occipital_inf |
|  | *v-occ* | Pole_occipital, G_oc-temp_med-Lingual, S_collat_transv_post, G_and_S_occipital_inf |
|  | *ctrl* | S_front_middle, G_front_middle, S_front_inf, S_front_sup |
| Glasser |  |  |
| Left | *d-occ* | V1 (1), V2 (4), V3 (5), V4 (6) |
|  | *v-occ* | V1 (1), V2 (4), V3 (5), V4 (6), PIT (22) |
|  | *ctrl* | 8Av (67), 8Ad (68), 8c (73), p9-46v (83), 46 (84), 9-46d (86) |
| Right | *d-occ* | V1 (1), V2 (4), V3 (5), V4 (6), LO1 (20), LO2 (21), V3CD (158) |
|  | *v-occ* | V1 (1), V2 (4), V3 (5), V4 (6) |
|  | *ctrl* | 8Av (67), 8Ad (68), 8c (73), p9-46v (83), 46 (84) |

**Table S2.** MNI coordinates of the boundaries of the average ROI. Boundaries for the average ROIs (d-occ and v-occ), and control region. Each coordinate represents the edge of the region in one of 6 directions. Coordinates in MNI space.

|  |  | ROI limits in MNI Neurosynth coordinates (https://neurosynth.org/) |  |  |  |  |  |
| --- | --- | --- | --- | --- | --- | --- | --- |
|  |  | superior | inferior | anterior | posterior | medial | lateral |
| Left | *d-occ* | -30, -93, 18 | -17, -101, -17 | -33, -82, 2 | -16, -105, -9 | -8, -104, 13 | -44, -92, 2 |
|  | *v-occ* | -19, -91, -5 | -27, -91, -21 | -33, -80, -14 | -9, -100, -10 | -8, -98, -11 | -48, -85, -11 |
|  | *ctrl* | -32, 33, 50 | -42, 34, 24 | -32, 43, 41 | -28, 19, 40 | -19, 29, 38 | -55, 23, 33 |
| Right | *d-occ* | 42, -81, 13 | 36, -91, -11 | 39, -75, 2 | 31, -100, -3 | 27, -92, 4 | 47, -84, 2 |
|  | *v-occ* | 14, -99, -7 | 30, -86, -19 | 30, -78, -17 | 18, -104, -7 | 9, -100, 1 | 42, -86, -16 |
|  | *ctrl* | 33, 23, 59 | 48, 25, 25 | 31, 39, 45 | 41, 15, 32 | 23, 30, 40 | 56, 25, 31 |

**Table S3.** Individual and mean R^2^ threshold. The R^2^ thresholds used to calculate the linear fits when quantifying the contentopic organization (see figure 4B). To ensure sufficient datapoints were included to perform the linear fits for our control region, we used a variable threshold (median). For our regions of interest, the threshold corresponded to an R^2^ cut-off of approximately 10 percent. For the control region the cut-off was an R^2^ of approximately 4 percent.

|  |  | **R2 threshold used (50% quantile)** |  |  |
| --- | --- | --- | --- | --- |
|  |  | **ctrl** | **occ** | **vent** |
| *sub1* | *left* | 0,042 | 0,132 | 0,103 |
|  | *right* | 0,038 | 0,113 | 0,114 |
| *sub2* | *left* | 0,038 | 0,113 | 0,073 |
|  | *right* | 0,036 | 0,086 | 0,106 |
| *sub3* | *left* | 0,044 | 0,136 | 0,120 |
|  | *right* | 0,049 | 0,122 | 0,123 |
| *sub4* | *left* | 0,032 | 0,103 | 0,107 |
|  | *right* | 0,033 | 0,067 | 0,094 |
| *sub5* | *left* | 0,043 | 0,091 | 0,116 |
|  | *right* | 0,048 | 0,088 | 0,092 |
| *sub6* | *left* | 0,048 | 0,104 | 0,107 |
|  | *right* | 0,041 | 0,086 | 0,101 |
| *sub7* | *left* | 0,038 | 0,133 | 0,139 |
|  | *right* | 0,040 | 0,102 | 0,132 |
| *sub8* | *left* | 0,040 | 0,101 | 0,126 |
|  | *right* | 0,035 | 0,119 | 0,101 |
| ***mean*** |  | 0,040 | 0,106 | 0,110 |

